# Enhanced functionalities of immune cells separated by microfludic lattice: assessment based on holotomography

**DOI:** 10.1101/2023.08.21.554226

**Authors:** Mahn Jae Lee, Byungyeon Kim, Dohyeon Lee, Geon Kim, Yoonjae Jung, Hee Sik Shin, Sungyong Choi, YongKeun Park

## Abstract

The isolation of white blood cells (WBCs) from whole blood constitutes a pivotal process for immunological studies, diagnosis of hematologic disorders, and the facilitation of immunotherapy. Despite the ubiquity of density gradient centrifugation in WBC isolation, its influence on WBC functionality remains inadequately understood. This research employs holotomography to explore the effects of two distinct WBC separation techniques, namely conventional centrifugation and microfluidic separation, on the functionality of the isolated cells. We utilize three-dimensional refractive index distribution and time-lapse dynamics to conduct an in-depth analysis of individual WBCs, focusing on their morphology, motility, and phagocytic capabilities. Our observations highlight that centrifugal processes negatively impacts WBC motility and phagocytic capacity, whereas microfluidic separation yields a more favorable outcome in preserving WBC functionality. These findings emphasize the potential of microfluidic separation techniques as a viable alternative to traditional centrifugation for WBC isolation, potentially enabling more precise analyses in immunology research and improving the accuracy of hematologic disorder diagnoses.

## 1. Introduction

Modifications in the morphology and function of white blood cell (WBC) subtypes act as diagnostic markers for a wide array of disorders, such as infections, cancer, and autoimmune diseases [1, 2]. Hence, the isolation and characterization of WBCs into morphological and functional subtypes have become a focal point of interest across the biological and medical disciplines, including cell phenotyping and immunotherapy [3, 4]. Cell analysis techniques such as flow cytometry, which evaluates cell count, activity, and morphology, have been developed and are in extensive use [5]. However, conventional separation methods like density gradient centrifugation present significant limitations. These include laborious procedures, potential alteration in immune cell phenotype, low purity and yield, and undesirable activation [6, 7]. In response to these challenges, microfluidic devices, which gently segregate WBCs based on size and morphology, have recently emerged as a viable alternative [8, 9].

Microscopy techniques, crucial for observing and analyzing cells, have conventionally relied on light microscopy, fluorescence microscopy, and electron microscopy [10-12]. While these methods have contributed to our understanding of immune cell structure and function, they come with certain constraints. Light microscopy, despite its ubiquitous use, provides limited resolution and often requires staining or labeling that can potentially disrupt the cell’s natural state or functions. Fluorescence microscopy, while allowing the visualization of specific cellular components, necessitates the use of fluorescent labels or markers which might jeopardize cell viability and function. Issues such as photobleaching and phototoxicity further limit the observation of live cells over extended periods [13]. More importantly, the use of exogeneous labeling agents prevent the use of cells for therapeutic applications. Electron microscopy, although it offers high-resolution imaging, requires intricate sample preparation and is incompatible with live cell imaging, thus posing significant challenges [14].

Quantitative Phase Imaging (QPI) has come forth as a robust solution, mitigating many limitations associated with traditional microscopy techniques [15-17]. As a label-free, non-invasive technique, QPI enables real-time observations of live cells and provides a quantitative analysis of parameters like cell morphology, concentration, mass, and volume. It utilizes the refractive index (RI) as an intrinsic quantitative imaging contrast [18]. The label-free and quantitative imaging capability of QPI an uncompromised study of cell imaging, making it the ideal method to authenticate the integrity of cells using microfluidic devices [19, 20].

In this research, we delve into the effects of different isolation methods on the functionality of individual live immune cells. We extracted polymorphonuclear leukocytes (PMNLs) utilizing two distinct enrichment methods: density gradient centrifugation and a microfluidic lattice chip, and compared the resultant morphological and functional modifications. Using holotomography (HT), a 3D QPI technique, we reconstructed the 3D RI tomogram of PMNLs to analyze their cellular morphology and functionality. By identifying the morphological and biophysical characteristics of individual immune cells and conducting time-lapse 3D RI imaging of live cells, we were able to investigate the dynamics of individual PMNLs. Our subsequent analysis of functional alterations illustrated the impacts of mechanical stress on immune cells. Our findings indicate that centrifugal force negatively impacts migration and phagocytosis capacities, whereas the microfluidic separator demonstrates potential in preserving WBC functionality.

## 2. Materials and Methods

### Fabrication and principle of microfluidic lattice

A microfluidic lattice separator was fabricated by standard photolithography and polydimethylsiloxane (PDMS) molding processes. The fabrication process of photoresist (PR) mold involved the procedures of cleaning a silicon wafer using a buffered oxide etchant, spin-coating of a PR layer to a thickness of 25 μm, baking of the PR mold at 100 °C, exposing it to ultraviolet light though a photomask, developing the PR using acetone, and post-development baking at 100 °C. The PDMS chip was then created by pouring a PDMS mixture of a base and a curing agent at 1:9 mass ratio and curing it at 100 °C for 10 min and 65 °C for 24 h. Once cured, the PDMS slab was peeled off from the mold and cut into individual chips. Inlet and outlet reservoir holes were drilled, and each chip was bonded to a glass slide after oxygen plasma treatment.

The PDMS chip integrates the principles of a microfluidic lattice separator and a pneumatic circuit to achieve pumpless WBC separation, as previously described [21, 22]. The microfluidic lattice consists of an array of main channels and side channels connecting them, which are designed to separate two blood cell types as WBCs flow along the main channels, and red blood cells (RBCs) exit through the separation channels (Fig. S1) [23]. Simultaneously, the lattice facilitates the washing of WBCs with a co-flowing buffer stream. This enables direct application of the separated WBCs to downstream analysis without the need for additional sample preparation, such as cell washing. To achieve pumpless operation of the separator, we used PDMS degassing. PDMS, being a gas-permeable material, can function as a built-in vacuum source for microfluidics upon degassing. Consequently, spontaneous separation can be achieved by filling the inlet reservoirs with blood and buffer solutions, while blocking the outlet reservoirs with a cover glass.

### Isolation of PMNLs

PMNLs were extracted from human peripheral blood samples employing two separation methods: density gradient centrifugation and microfluidic separation. The density gradient method selectively segregates cells falling within a specific mass density range, whereas the microfluidic device separates large WBCs from smaller red blood cells (RBCs) and blood plasma.

To isolate PMNLs using conventional centrifugation, we prepared a layered mix of density gradient media (Histopaque 1119, 1077), over which we delicately layered a solution of whole blood and phosphate-buffered saline (PBS) diluted with 2% Fetal Bovine Serum (FBS). Post-centrifugation at 800 g for 30 minutes without brake, granulocytes were identified in the median layer of the gradient media. The retrieved granulocytes were twice rinsed with 10 mL of PBS and centrifuged for 10 minutes at 200 g. We then resuspended the resultant cell pellet in 5 mL of PBS.

The microfluidic devices were utilized to separate WBCs containing PMNLs. The microfluidic devices were primed by infusing PBS with 0.1% BSA post-degassing of the devices. Once the microchannels were completely filled with the washing buffer, we introduced whole blood. The microfluidic lattice successfully separated WBCs from smaller red blood cells and blood plasma. A collection period of 15 min sufficed to gather an adequate number of WBCs for our ensuing experiments.

The sorted cells were subsequently transferred onto a TomoDish (Tomocube Inc., Republic of Korea) using a #1.5H coverslip (Paul Marienfeld GmbH & Co. KG, Germany) in preparation for acquiring the refractive index (RI) tomograms.

This study and all experimental protocols were approved by the KAIST Institutional Review Board (IRB project number: 2012-0128).

### Holotomography

To capture RI tomograms of live neutrophils, we employed an HT system (HT-2H; Tomocube Inc., Republic of Korea). This HT system functions based on Mach-Zehnder interferometry, using a coherent light source with a wavelength of 532 nm. Multiple 2D holograms, captured at different illumination angles modulated by exploiting a digital micromirror device [24, 25], are used to reconstruct the 3D RI tomogram of a cell, employing the Fourier diffraction theorem and a Rytov approximation for gradually varying RI [26]. Precise control over the illumination angle impinging on the sample is achieved through a digital micro-mirror device.

Despite these measures, due to the finite numerical apertures of both the condenser and objective lenses with the numerical apertures of 1.2, the reconstructed RI tomogram is subject to axial elongation, a phenomenon known as the ‘missing cone problem.’ We employed a regularization method predicated on a non-negativity constraint to address this issue. The theoretical resolution of the HT system is determined to be 156 nm and 360 nm for the lateral and axial directions, respectively [27]. Further details regarding the principle, instrumentation, and reconstruction code of HT are available in the referenced sources [27-29].

### Phagocytic functionality assay

The functionality of the purified neutrophils was assessed by evaluating their phagocytic capacity under microscopic observation. We employed fluorescently labeled E. coli (Alexa Fluor 594) as a functional probe. The E. coli stock was diluted to a concentration of 200 μg/ml and allowed to incubate for an hour at 37°C. We then exposed the purified neutrophils to this diluted mixture.

Following this, the cells were transferred to a plate (TomoDish; Tomocube Inc., Republic of Korea) and incubated for an additional hour at 37°C. Phagocytic neutrophils create a vesicle to bind to the bioparticles. To amplify the differentiation capability in the images, we employed a Hoechst stain. Thereafter, phagocytic uptake was examined using a combined fluorescence and HT system over a 30-minute observation period.

## 3. Results

### Holotomography of PMNLs

We extracted WBCs from whole blood using two distinct methodologies (Fig. 1A; Refer to Materials and Methods). In contrast to the centrifugation method, which required over an hour to acquire our target cells, the microfluidic chip accomplished cell separation in under 20 minutes. Following the sample preparation using both methods, we obtained 3D RI tomograms of the PMNLs using HT (Fig. 1B). HT, the optical analogous to X-ray computed tomography (CT), enabled the precise and high-resolution reconstruction of unlabeled live cells and tissues [16, 26, 30]. Due to its label-free and quantitative 3D imaging capability, HT has recently been utilized for the study of flow cytometry [19, 20], cell biology [31-33], immunology [34, 35], microbiology [36, 37], biomolecular condensates [31, 32], drug screening [38, 39], and regenerative medicine [40, 41].

**Fig 1.**
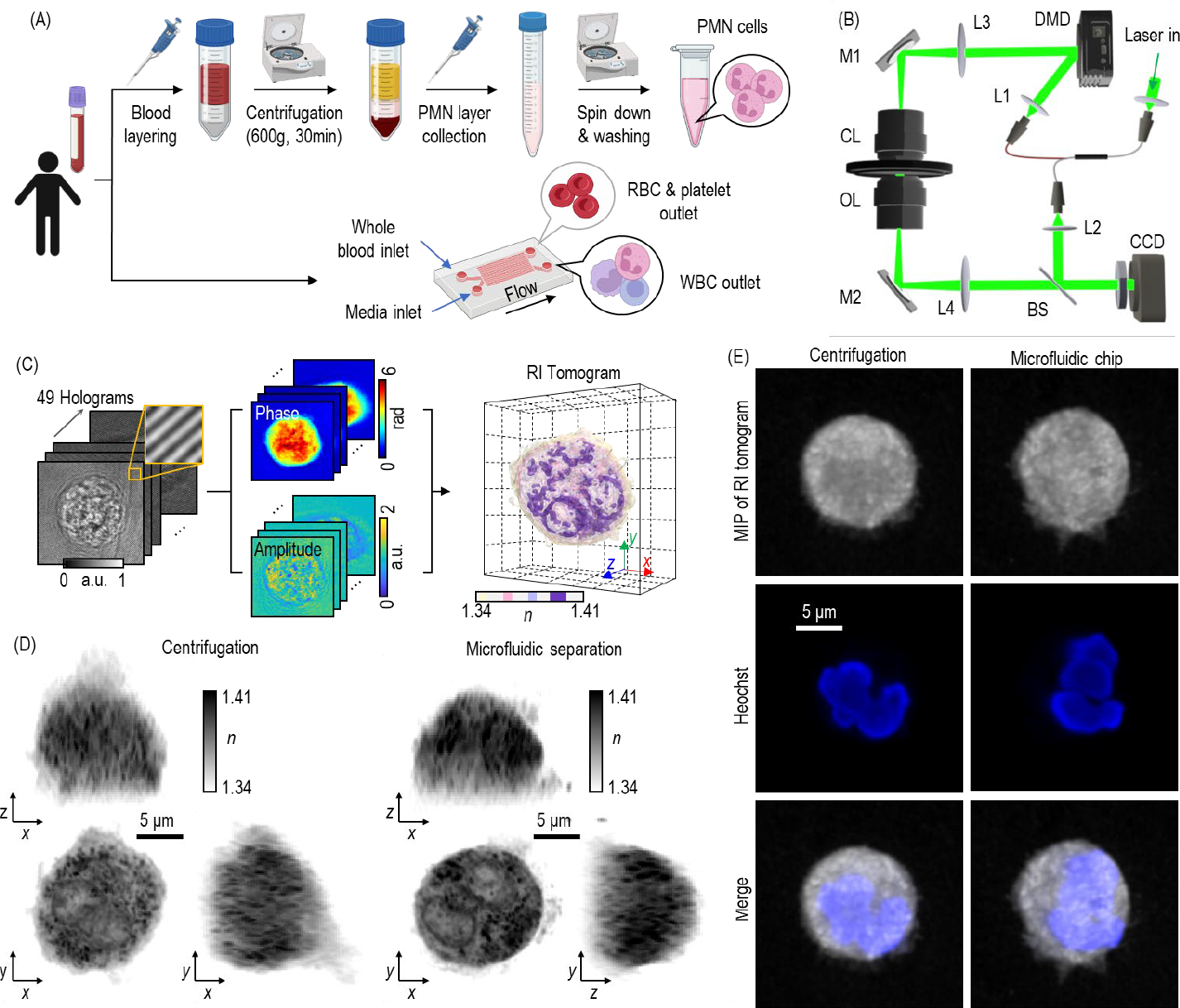
Experimental steps to compare proposed white blood cell isolation techniques. (A) White blood cells are extracted from whole blood samples employing two distinct methodologies. (B) Label-free Refractive Index (RI) tomography documents the condition of each isolated PMNL. (C) Reconstruction of the 3D RI tomogram using 49 holograms, with the interference pattern depicted in the orange magnified box. (D) Display of representative images of the reconstructed RI tomograms, including maximum contrast projections to three orthogonal planes. (E) Refractive index tomograms synergistically supplement information provided by three-dimensional fluorescence images.

In brief, the hologram of the scattered beam passing through the sample was acquired using 49 distinct holograms, enabling the reconstruction of the 3D RI tomograms (Fig. 1C). The reconstructed RI tomograms exhibited an RI value range from 1.34 to 1.41, aligning with previously reported findings [42]. The representative images display PMNLs composed of multi-lobulated nuclei and multiple intracellular granules, which demonstrated a high RI (Fig. 1D). To ensure our study focused solely on PMNLs, we used a fluorescence-based validation, where the characteristic nuclear morphology was inspected to exclude other types of WBCs (Fig. 1E).

### Comparison of biophysical and morphological characteristics

The two isolation techniques, namely, centrifugation and microfluidics, were compared based on the cellular attributes obtained from HT in terms of biophysical and morphological characteristics. We began with segmenting cells utilizing Otsu’s method (Fig 2A) [43]. The segmentation with RI values served to evaluate parameters such as volume, protein density, dry mass, solidity, and eccentricity (Fig 2B) from the measured 3D RI tomogram of individual cells. There is a linear relationship between RI and dry mass concentration: *Δn* = *α·C*, where *Δn* is the RI difference between cells and surrounding media and *C* represents the cell’s dry mass concentration, was used to estimate the dry mass of a single cell [44]. The RI increment α was established at 0.185 μm^3^/pg [45]. The resulting statistics revealed no significant difference in dry mass between the two isolation methods. Solidity, a measure of the convexity or concavity of the cell’s shape, and eccentricity, a measure of the cell’s elongation, provided insight into the cellular form. Generally, cells with irregular shapes exhibit lower solidity and higher eccentricity.

**Fig 2.**
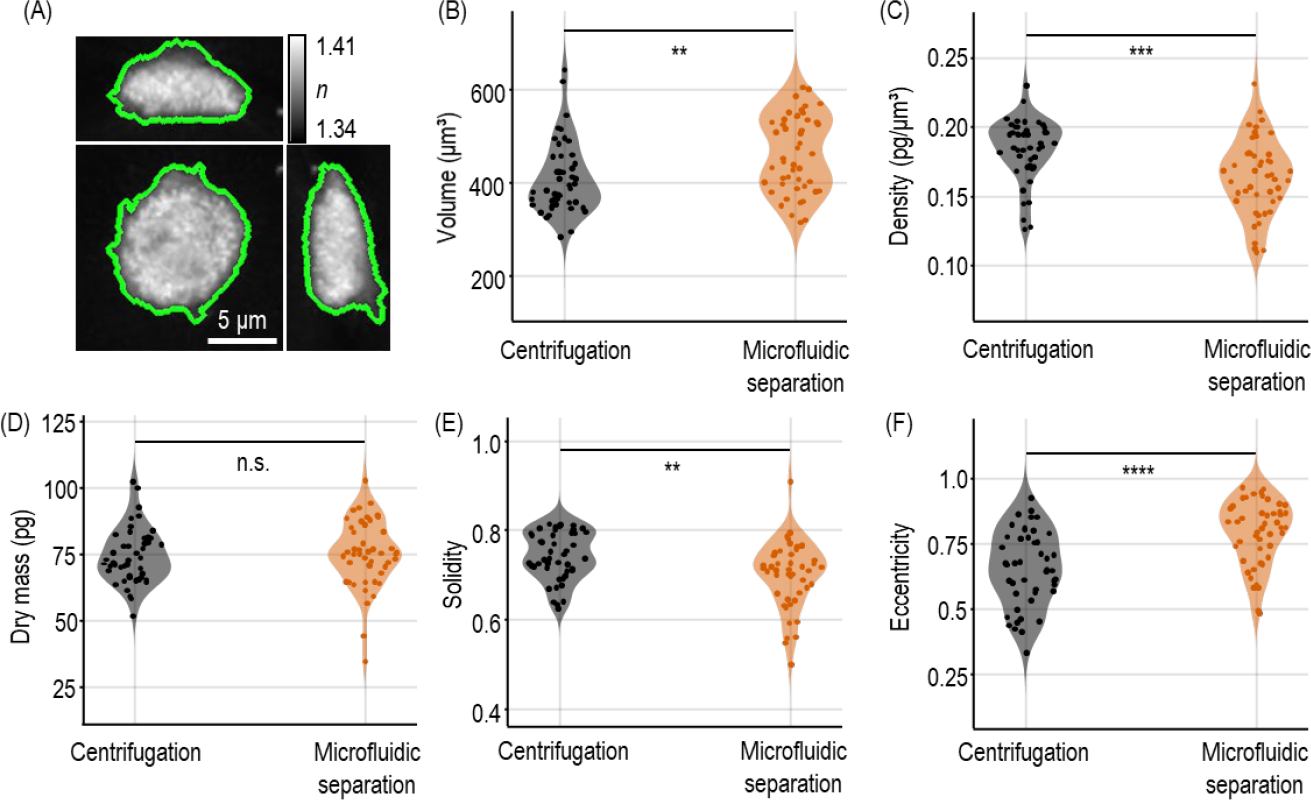
Statistical Examination of morphological and biophysical features. (A) Maximum contrast projections of PMNLs, shown representatively. Otsu’s method-based established algorithm segregates PMNLs from the background (illustrated by the green line). (B) Morphological analysis reveals distinct behaviors of PMNLs dependent on the isolation technique employed. Distributions were statistically evaluated using the two-sided unpaired t-test. Significance levels are indicated as follows: \*\**p*<0.01, \*\*\**p*<0.001, \*\*\*\**p*<0.0001, while n.s. denotes non-significant differences with p>0.05.

Notably, the phenotypic parameters indicated significant variations in PMNLs contingent upon the isolation techniques. PMNLs separated by centrifugation exhibited higher dry mass compared to those separated by the microfluidic chip. This discrepancy may stem from the distinct operating principles of the two methods; the microfluidic chip primarily separates particles based on size rather than density [46]. Irrespective of the isolation method, the dry mass remained constant. The solidity and eccentricity parameters suggested that PMNLs obtained through the microfluidic chip exhibited more irregular forms. This finding aligns with previous reports suggesting that centrifugation causes pseudopod retraction [47].

### Analysis of cellular motility

The migratory capacity of PMNLs plays a pivotal role in immune defense. To evaluate this capacity, we monitored the movement of individual cells every second using holotomography (HT) (Fig. 3A). Subsequently, these cell trajectories were analyzed using an anomalous diffusion model to quantify the dynamic behavior into discernible values (Fig. 3B) [48]. The anomalous diffusion model evaluates cellular viability and mobility, represented by the diffusion exponent (*α* in Fig. 3B) and the diffusion coefficient (K in Fig. 3B).

**Fig 3.**
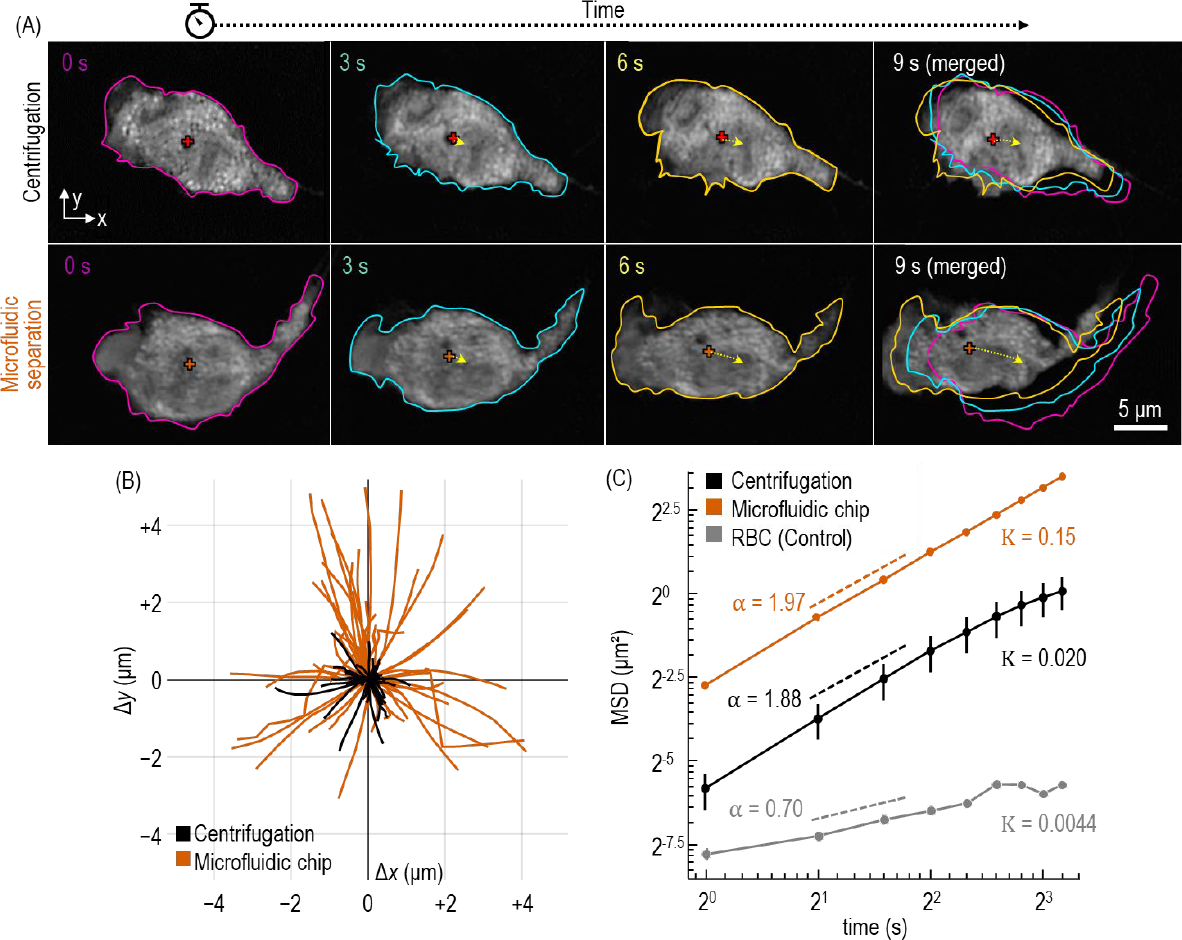
Analysis of cellular dynamics. (A) Single PMNL cells are continuously tracked and visualized using RI tomograms at three-second intervals. The cellular boundary is represented by different colors for each time frame. (B) A diffusion model is utilized to quantify the mobility of PMNLs. Error bars represent standard deviations. MSD denotes Mean Squared Displacement.

The fluctuation in the cellular boundary reflects the active crawling behavior of stress-free PMNLs (Fig. 3A). These PMNLs exhibited a heightened level of activity, frequently extending and retracting their pseudopods. This active movement is a characteristic behavior of stress-free PMNLs, as demonstrated in the trajectory map (Fig. 3B). We validated our analysis using linear regression with the anomalous diffusion model. Red blood cells were also analyzed as a control group. The data fitting results suggested that centrifugal force diminishes the locomotive capacity of PMNLs. This observation aligns with our previous morphological analysis regarding solidity and eccentricity, as a retracted pseudopod can influence locomotion capacity [49]. The results suggest that the microfluidic chip facilitates the separation of PMNLs under minimal stress, thereby preserving their active locomotion capacity.

### Phagocytosis capacity analysis

We aimed to assess the phagocytic capability of PMNLs isolated using two distinct methods by co-culturing them with fluorescently labeled *E. coli* bioparticles (Fig. 4A). The phagocytic capacity was quantified by measuring the fluorescence intensity. RI tomograms revealed that phagocytosed bioparticles possess a relatively high RI value (Fig. 4B). Our data suggested that PMNLs derived via microfluidic chip separation demonstrated greater fluorescence intensities compared to those isolated through the centrifugation method (Fig. 4C). Given the assumption that the average fluorescence intensities of *E. coli* bioparticles are consistent, the higher intensities imply an increased number of phagocytosed bioparticles. To determine the quantity of phagocytosed bioparticles, we utilized fluorescent signal thresholds for segmentation and estimated their dry mass using RI values. Our results indicated a higher dry mass of intracellular *E. coli* bioparticles in cells separated by the microfluidic chip, compared to those obtained through centrifugation. There was no significant difference in the mean RI of intracellular *E. coli* bioparticles between the two groups, suggesting that the composition of the phagocytosed bioparticles remained unchanged. The elevated dry mass of intracellular *E. coli* bioparticles indicates that the microfluidic technique preserves the phagocytic function of PMNLs more effectively than centrifugation.

**Fig 4.**
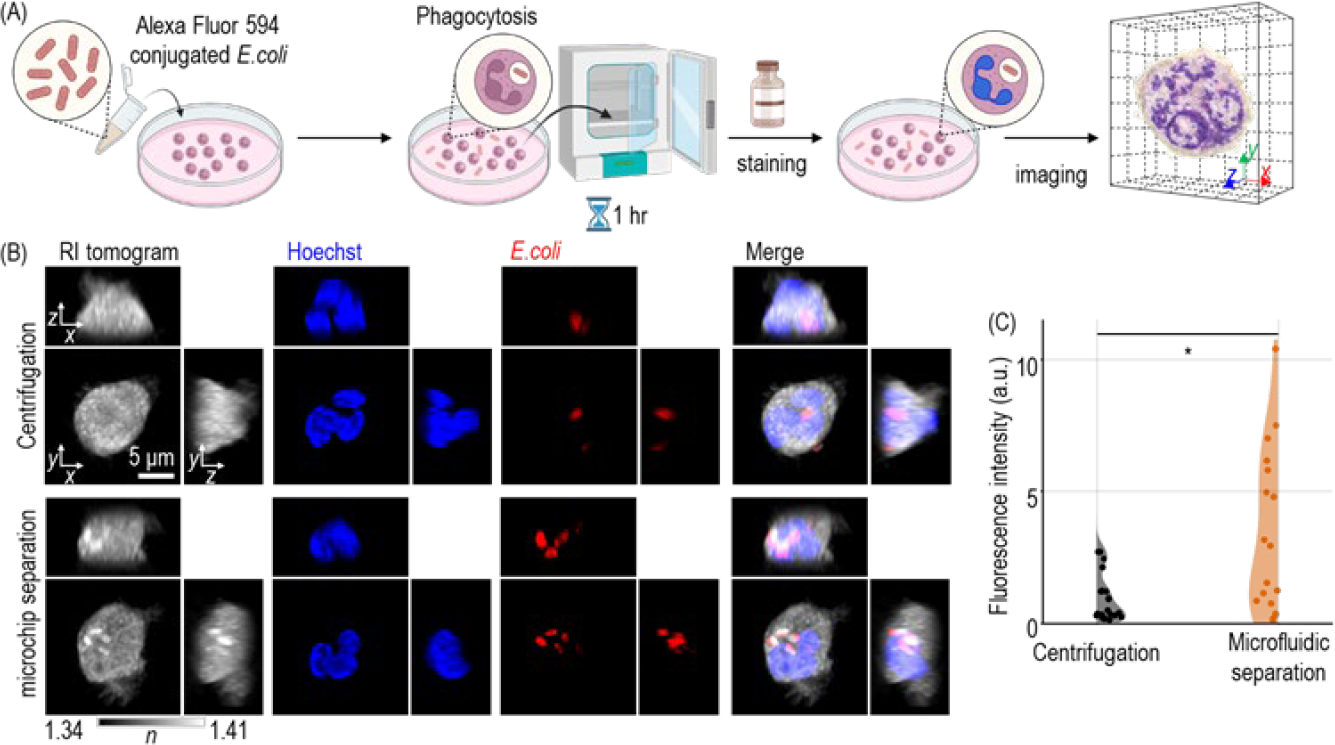
Phagocytic functionality assay. (A) The phagocytosis assay quantifies the immunological response of PMNLs. (B) Combined RI tomogram and fluorescence images enable the counting of only fluorescently-conjugated E. coli within each PMNL. (C) PMNLs isolated using the microfluidic chip demonstrate higher phagocytic capacity compared to those isolated via the conventional centrifugation method. Distributions were statistically compared using a two-sided unpaired t-test. \**p*<0.05.”

## 4. Discussion

Centrifugation, a conventional technique for cell separation, enrichment, and washing, has been widely used due to its limited effect on the viability of pliable and elastic biological specimens, such as cells. However, researchers have often overlooked the morphological and biophysical changes caused by centrifugation, which may be crucial in areas like clinical diagnostics that depend on cell morphology observation. Our study uncovered significant alterations in cellular morphology and biophysical properties as a result of centrifugation, including decreased motility and phagocytic capacity. Using a microfluidic chip can mitigate these centrifugal influences on cells. Moreover, the microfluidic chip better simulates the architecture of human blood vessels, offering a more accurate representation of in situ cellular dynamics.

One possible explanation for the morphological change is cell dehydration caused by centrifugation. Given that the RI is directly proportional to protein concentration, and there is no significant difference in the dry mass of cells between the two groups, the increased intracellular protein concentration in centrifuged cells suggests a reduction in intracellular volume. As the primary constituent of cells is water, this decline in volume can be largely attributed to dehydration.

Centrifugal force-induced cytoskeletal remodeling could also impact cell morphology and functionality [50]. Most cells possess a filamentous actin (F-actin) cytoskeleton responsible for bearing the majority of mechanical stress and maintaining cell shape. Mechanical forces can induce F-actin reconfiguration, leading to alterations in cellular morphology. The cytoskeleton is involved in leukocyte migration, immunological synapse formation, and phagosome formation, aligning with our statistical analysis results [51]. Further investigation into intracellular cytoskeletal changes might provide more insights into the impact of centrifugal force on cells. Moreover, employing machine-learning-based image inference could be beneficial for virtually generating molecular-specific imaging without the need of introducing exogenous labeling agents [52].

The results shown in this work suggest the development of a label-free biomarker to indicate cellular functionality. Exploiting both static and dynamic RI tomograms of individual WBCs has the potential to create a virtual biomarker correlating to cellular functionality. Static RI tomograms provide intrinsic biophysical information about cellular structures and morphologies, including their dry mass, protein concentration, and shape features. These parameters can be associated with various cell functions, such as cell type and maturity, signaling pathways, or disease states. On the other hand, dynamic RI tomograms can give insights into the temporal behavior of cells, illustrating the changes in intracellular structures and cell morphologies over time. These could be linked to active cellular processes such as migration, proliferation, differentiation, and response to environmental changes. By integrating both static and dynamic information, a comprehensive and rich biomarker could be constructed. This biomarker could be a robust indicator of the WBC’s functionality, and it may be applicable to a wide range of studies, from basic biological research to clinical diagnostics and prognostics, thereby opening a new window to understand the complex cellular dynamics in a more refined manner.

This study does also have a few limitations. First, this study focused on PMNLs, and the conclusions may not necessarily apply to other WBC types. Each type of WBCs has unique biophysical characteristics and functions, and the effects of the isolation method could differ among them. Future studies could focus on expanding the research to other cell types. Second, current microfluidic device is small and may not be practical for handling large volumes of blood, which might be necessary in clinical settings. Scaling up the microfluidic device without compromising its effectiveness would be a challenge. Third, although the study effectively measures basic cellular functions like migration and phagocytosis, it does not directly address more complex and subtle biological processes, such as signal metabolism, transduction or gene expression changes. The use of fluorescently labelled *E. coli* bioparticles to evaluate phagocytic function is an artificial condition that may not fully reflect the cells’ behavior in the human body. These limitations could potentially be addressed in future work. For instance, a more diverse set of WBCs could be tested to generalize the findings. The microfluidic device could be redesigned to handle larger volumes. More sophisticated techniques, potentially involving machine learning or other computational methods, could be developed to analyze more complex biological processes. Additionally, more physiologically relevant assays could be developed to better assess cellular functionality in conditions that mimic the human body more closely.

In conclusion, our methodology enabled the label-free evaluation of cellular morphology and functionality. The microfluidic chip facilitated rapid enrichment of WBCs, substantially reducing sample preparation time. Time-lapse imaging at high temporal resolution allowed for the measurement and differentiation of phagocyte dynamics, distinguishing them from anomalous diffusion and suggesting a novel quantitative parameter for migration capacity. Furthermore, correlative imaging with fluorescent bioparticles allowed for the examination and comparison of phagocytic capacity based on different separation techniques. Our findings showed that centrifugal force negatively impacts both migration and phagocytosis capacities of phagocytes, and these effects can be counteracted by using a microfluidic chip. We believe that our study introduces a new approach that minimizes cellular deformation for image analysis.

## Data availability

Data underlying the results presented in this paper are not publicly available at this time but may be obtained from the authors upon reasonable request.

