## Supplementary Figure 1 for "Enhanced functionalities of immune cells separated by microfludic lattice: assessment based on holotomography"

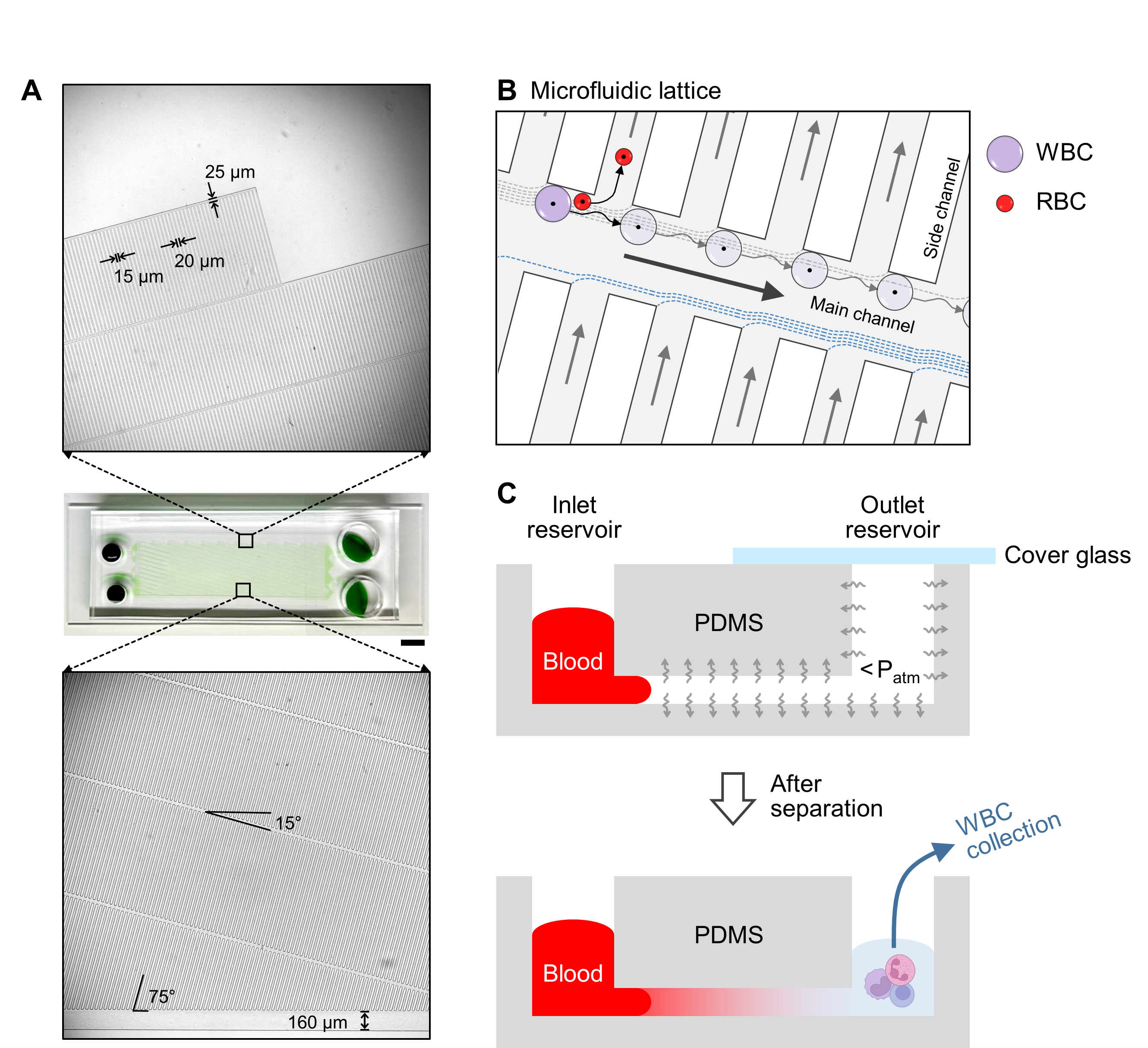

**Fig. S1.** (A) Photography of a microfluidic lattice separator with enlarged insets showing main and side channels in details. Scale bar, 5 mm. (B) Schematic illustration of microfluidic lattice-based blood cell separation process. (C) Schematic illustration of autonomous microfluidic pumping based on PDMS degassing and subsequent collection of sorted white blood cells after microfluidic separation.
